# Metabolomic and proteomic analyses of a quiescent *Escherichia coli* cell factory reveal the mechanisms behind its production efficiency

**DOI:** 10.1101/082305

**Authors:** Nicholas M. Thomson, Tomokazu Shirai, Marco Chiapello, Akihiko Kondo, Krishnan J. Mukherjee, Easan Sivaniah, David K. Summers, Keiji Numata

## Abstract

Quiescent (Q-Cell) *Escherichia coli* cultures can be created by using the signalling molecule indole to halt cell division of an *hns* mutant strain. This uncouples metabolism from cell growth and allows for more efficient use of carbon feedstocks. However, the reason for the increased productivity of cells in this state was previously unknown. We show here that Q-cells can maintain metabolic activity in the absence of growth for up to 24 h, leading to four times greater per-cell productivity of a model metabolite, 3-hydroxybutyrate (3HB), than a control. Metabolomic data show that by disrupting the proton-motive force, indole interrupts the tricarboxylic acid cycle, leading to the accumulation of metabolites in the glycolysis pathway that are excellent starting points for high-value chemical production. By comparing protein expression patterns between wild-type and Q-cell cultures we show that Q-cells overexpress stress response proteins, which prime them to tolerate the metabolic imbalances incurred through indole addition. Quiescent cultures produced half the cell biomass of control cultures lacking indole, but were still able to produce 39.4 g.L^-1^ of 3HB compared to 18.6 g.L^-1^ in the control. Therefore, Q-cells have high potential as a platform technology for the efficient production of a wide range of commodity and high value chemicals.

## 2 Introduction

In *E. coli*, carbon flux through the central carbon metabolism pathways of glycolysis, the pentose phosphate pathway (PPP) and the tricarboxylic acid cycle (TCA) is at a maximum during exponential growth (Meier et al., 2011). This offers abundant opportunities for re-routing of metabolic pathways for the commercial scale production of high value or commodity chemicals from cheap, renewable feedstocks (Chen et al., 2013; Li, 2011; Murphy, 2012; Yu et al., 2011). Consequently, *E. coli*-based bioproduction processes usually aim to extend the exponential phase for as long as possible before nutrient limitation, oxygen deprivation or the accumulation of toxic by-products inhibit further carbon metabolism and cell growth (Choi et al., 2006; Lee, 1996; Singh et al., 2012). The consequential accumulation of biomass (essentially a waste product) requires the diversion of feedstock for maintenance of cell function and production of essential macromolecules (Van Bodegom, 2007).

Alternatives to high cell density growth are used in certain circumstances, including continuous production using a chemostat (Hoskisson and Hobbs, 2005; Hua et al., 2004) and the use of ‘resting cells’ in an osmotically-balanced but nutrient-limited buffer such as phosphate buffered saline (Cha et al., 1999; Ghazi et al., 1983). However, these approaches are technically more challenging and suffer from problems including vulnerability to contamination in chemostats and limited viable lifespans of resting cells.

Quiescence is achieved by addition of 2.5 – 3.0 mM indole to cultures of *E. coli* W3110 carrying a stop codon after the 93^rd^ codon of the *hns* gene (*hns*Δ93). This mutation causes the production of a truncated Histone-like Nucleoid Structuring protein (H-NS). After indole is added, metabolic activity continues and production of plasmid-encoded proteins is increased, but only in *hns* mutants (Chen et al., 2015; Mukherjee et al., 2004; Rowe and Summers, 1999). Indole is a well-studied chemical signal in over 85 species of bacteria (Lee and Lee, 2010). It is a proton ionophore and has been shown to reduce the proton-motive force (PMF) of *E. coli* by allowing protons to return to the cytoplasm after their expulsion to the periplasmic space during oxidative phosphorylation (Chimerel et al., 2013). One effect of reducing the PMF is to prevent the formation of the FtsZ ring, which is a prerequisite for cell division. Therefore, at suitable concentrations indole is able to prevent *E. coli* cell division (Chimerel et al., 2012).

Here, we investigated the changes in central carbon metabolism and protein expression that are induced by indole addition in wild-type and Q-cell (hnsΔ93) cultures under fed-batch conditions. Taking into account the results of these experiments, we designed a simple fermentation strategy for improved production of 3HB.

## 3 Results

We conducted a time-course study by growing the wild-type or *hns*Δ93 mutant strain of *E. coli* W3110 in 3 L aerobic, fed-batch fermentations with or without the addition of indole. The *hns*Δ93 cultures grew slower than the wild-type (Fig. 1a). Therefore, we normalized the early culture conditions by inoculating into modified mineral salts medium (M9GYT) containing 0.4% glucose and growing until the initial carbon supply was exhausted. At this stage, every culture reached the same optical density at 600 nm (OD_600_; overall average 6.69, standard deviation 0.48). The nutrient feed was then started and the time-course was begun 30 min later by addition of indole (final concentration of 3 mM from a 1 M stock dissolved in ethanol) or an equivalent volume of pure ethanol. Only *hns*Δ93 cultures entered quiescence, whereas wild-type growth was inhibited but not stopped by indole.

**Figure. 1:**
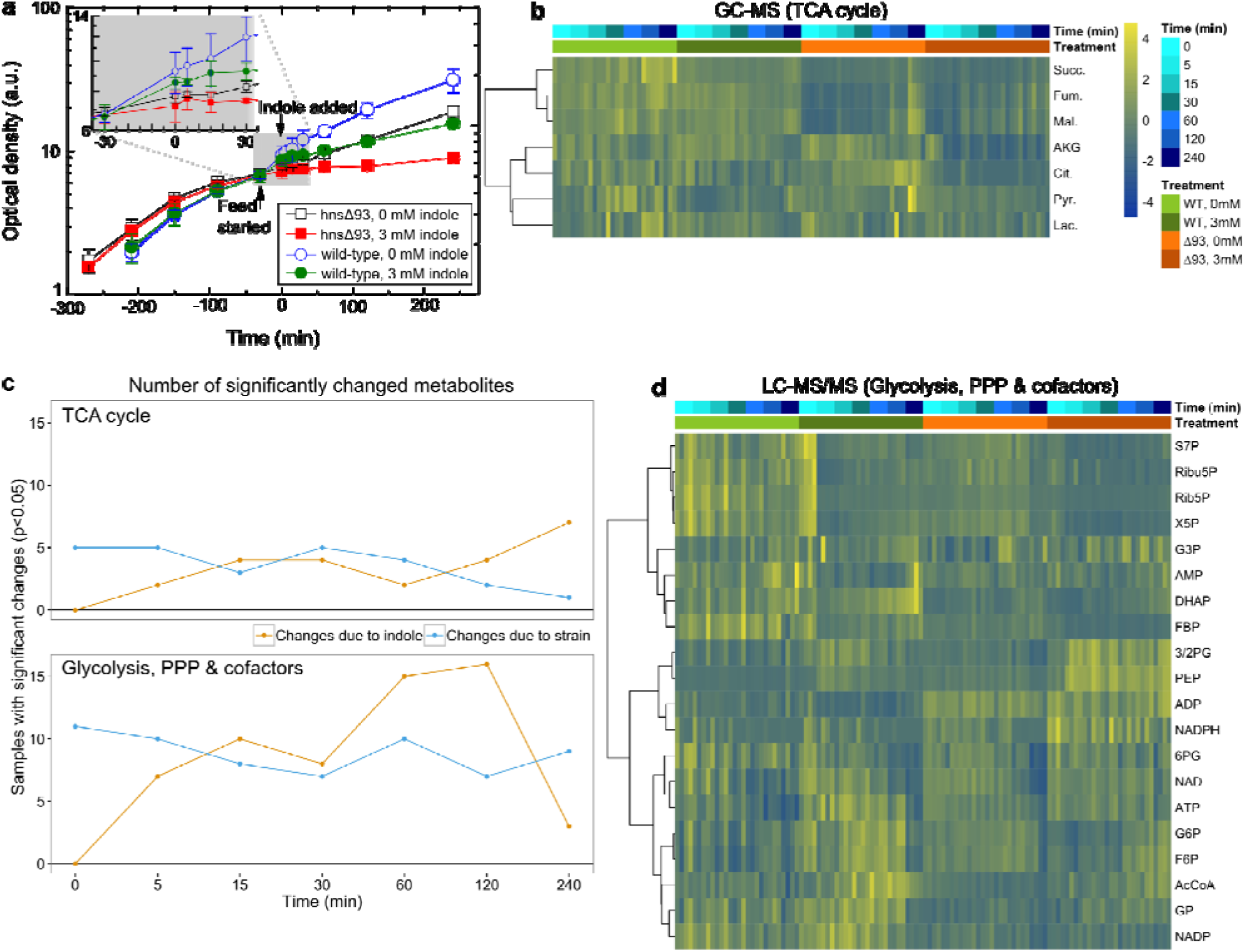
Growth and changes in metabolite concentration. **a**, Growth curves of each strain during the time-course experiment. The inset graph shows a magnified area of the same data from the time feeding was started until 30 mins into the time-course. Error bars represent the standard deviations (n = 4). **b**, Heat map of concentration changes in intermediates of the TCA cycle plus pyruvate and lactate, analyzed by GC-MS. **c**, Summary of the individual ANOVA analyses for each time-point showing the increasing importance of indole-induced concentration changes over time. **d**, Heat map of concentration changes in cofactors and intermediates of glycolysis and PPP analyzed by LC-MS/MS. Abbreviations for metabolite names: S7P, sedoheptulose-7-phosphate; Ribu5P, ribulose-5-phosphate; Rib5P, ribose-5-phosphate; X5P, xylulose-5-phosphate; G3P, glyceraldehyde-3-phosphate; AMP, adenosine monophosphate; DHAP, dihydroxyacetone phosphate; FBP, fructose-1,6-bisphosphate; 3/2PG, 3-phosphoglycerate & 2-phosphoglycerate; PEP, phosphoenolpyruvate; ADP, adenosine diphosphate; 6PG, 6-phosphogluconate; ATP, adenosine triphosphate; G6P, glucose-6-phosphate; F6P, fructose-6-phosphate; AcCoA, acetyl-Coenzyme A; GP, glucose-1-phosphate; NAD, oxidized nicotinamide adenine dinucleotide (NAD); NADH, reduced (NAD); NADP, oxidized NAD phosphate (NADP); NADPH, reduced NADP; Succ., succinate; Fum., fumarate; Mal., malate; AKG, α-ketoglutarate; Cit., citrate; Pyr., pyruvate; Lac., lactate.

We analyzed 22 cofactors and intermediates of glycolysis and PPP using liquid chromatography – tandem mass spectrometry (LC–MS/MS) and 8 organic acids (pyruvate, lactate and intermediates of TCA) by gas chromatography – mass spectrometry (GC–MS). Glucose-ADP, isocitrate and NADH were removed from the analysis due to very low abundances. The quantitative (GC–MS and HPLC) and relative ratio (LC–MS/MS) concentrations for the remaining 27 species were normalized against the OD_600_ of the cultures at each time point. Results using each technology were then analyzed in parallel, since direct comparisons between the two concentration reporting modes were not possible.

### 3.1 The metabolic effects of indole are strongest during exponential growth

Metabolites tended to cluster with neighboring intermediates of their respective pathways (Fig. 1b,d), showing that any metabolic changes tended to affect whole pathways more than individual intermediates. Individual principal component analyses (PCA) for each time-point of the LC-MS/MS data revealed a dynamic interaction between the influences of strain differences and indole (Fig. 2, Supplementary Fig. 1). Before indole addition, the samples were relatively closely clustered with some separation by strain along principal component (PC) 1 (accounting for 56% of the observed difference). Within 5 min of indole addition, indole treatment became the dominant clustering factor, while PC1 became less important (38.3% of observed differences). The influence of indole became more pronounced over the following 60 min, but by 240 min the samples tended to cluster more closely again, with some separation by strain. Untreated culture density was already high by this time and growth may have begun to slow. This suggests that the quiescent metabolic state resembles the late-logarithmic or early stationary phase of batch cultures. The GC-MS data revealed a similar trend for organic acids (Supplementary Fig. 2).

**Figure 2:**
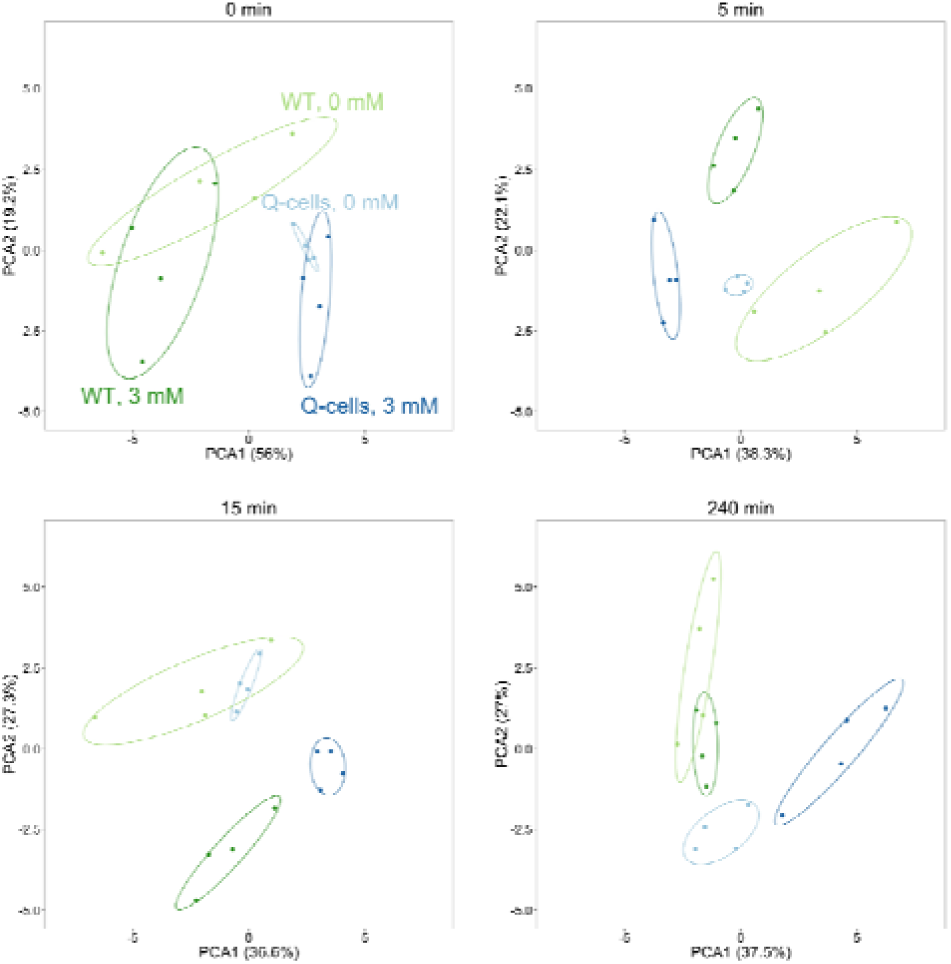
Individual principal component analyses of four time points for the metabolites analyzed by LC-MS/MS. These time-points were chosen to demonstrate the rapid (< 5 min) but reversible (< 240 min) response to indole addition, which outweighed the differences between the wild-type and *hns*Δ93 strains. Points represent individual fermentations (n = 4) and ellipses represent 95% confidence intervals. The full series of PCA analyses for LC-MS/MS and GC-MS data are shown in Supplementary Fig. 1 and 2.

We also observed the temporally changing influence of indole by conducting individual ANOVA tests for each time-point (Fig. 1c). The number of metabolites with a significant change due to strain remained fairly constant throughout the time-course, whereas indole-associated changes increased over 120 min before falling again by 240 min.

### 3.2 Indole provides an increased pool of available metabolites from glycolysis in hnsΔ93 cells

To identify the metabolites with the most significant changes in concentration during quiescence, we compared the time-courses of each metabolite between the control (wild-type, 0 mM indole) and quiescent cultures, and ranked each metabolite by the variation in time-course profile using Multivariate Empirical Bayes Analysis for time-course data (MEBA) (Tai and Speed, 2006). The individual graphs of concentration versus time for each metabolite and cofactor are provided in Supplementary Fig. 3 – 6. Intermediates of PPP were unchanged or reduced in concentration over time under either condition (Table 1). However, glycolysis intermediates were present at higher concentrations in quiescent cells than in the control. We noted particularly significant increases in phosphoenolpyruvate (PEP), 3-phosphoglycerate and 2-PG (3/2-PG), and acetyl-Coenzyme A (Ac-CoA), of which the latter two increased over time in quiescent cells but decreased in the control.

**Table 1:** Most significantly changed metabolites during the time-course experiment.

| Metabolite | Pathway | Fold change<br>in WT, 0 mM | Fold change<br>in hnsΔ93, 3 mM | Hotelling T<sup>2</sup> |
| --- | --- | --- | --- | --- |
| *LC-MS/MS* |  |  |  |  |
| Phosphoenolpyruvate | Glycolysis | 1.13 | 2.91 | 87.62 |
| Sedoheptulose-7-P | PPP | -1.03 | -1.23 | 58.72 |
| NAD | Cofactor | -1.48 | -1.28 | 53.53 |
| Fructose-1,6-BP | Glycolysis | -1.65 | -1.30 | 50.44 |
| 3/2-phosphoglycerate | Glycolysis | -1.01 | 1.53 | 48.97 |
| ADP | Cofactor | -1.28 | 1.09 | 48.66 |
| Acetyl-CoA | Glycolysis/TCA | -1.27 | 1.41 | 42.27 |
| Ribose-5-P | PPP | -2.56 | -1.38 | 35.69 |
| Xylulose-5-P | PPP | -1.76 | -1.70 | 34.58 |
| Ribulose-5-P | PPP | -1.35 | -1.76 | 33.05 |
| *GC-MS* |  |  |  |  |
| α-ketoglutarate | TCA | -1.13 | -1.52 | 95.42 |
| Malate | TCA | 1.03 | 2.00 | 73.18 |
| Succinate | TCA | 1.72 | 2.06 | 72.79 |
| Fumarate | TCA | 1.16 | 1.74 | 70.56 |
| Citrate | TCA | 1.23 | -2.81 | 64.18 |
| Pyruvate | Glycolysis/TCA | 1.64 | -1.63 | 49.01 |
| Lactate | Other | 1.57 | -1.51 | 42.51 |

Both strains had identical flux ratios through glycolysis, PPP and the Entner-Doudoroff pathway in ^13^C-labelled chemostat experiments without indole (Supplementary Table 1). Due to its effect on growth, we could not investigate the changes in metabolic flux caused by indole. However Fig. 1b,d and Table 1 both suggest that indole causes a non-H-NS dependent re-routing of metabolic flux away from the PPP, which results in a build-up of intermediates in the glycolysis pathway.

### 3.3 Indole inhibits the TCA cycle

MEBA also revealed an effect of quiescence on the TCA cycle (Table 1). In control cultures, TCA intermediates slowly increase in concentration during growth. However, in quiescent cultures pyruvate and acids from the first half of TCA reduced in concentration, whereas those in the second half increased above the levels of the control. With the simultaneous increase in Ac-CoA, this suggests indole inhibits the TCA cycle, possibly by preventing the conversion of Ac-CoA to citrate.

An alternative explanation could be that the glyoxylate cycle is stimulated in the presence of indole (Fig. 3). The glyoxylate cycle is an anabolic pathway that allows growth on simple carbon sources such as acetate in the absence of glucose, relying on isocitrate lyase (AceA) and two isozymes of malate synthase (AceB and GlcB) (Cronan, Jr. and Laporte, 2006; Maharjan et al., 2005). HPLC analysis of culture supernatants showed that acetate was the only organic acid secreted by the cells, and was produced at approximately 30% higher specific concentration by *hns*Δ93 than by wild-type cells. Since acetate stimulates the glyoxylate cycle (Maloy and Nunn, 1982; Ornston and Ornston, 1969) it is plausible that higher concentrations of acetate might result in a higher flux through the glyoxylate cycle even when glucose is present. Using quantitative real-time PCR (qRT-PCR) we measured *aceB* and *glcB* expression levels 1.6-fold (p < 0.01) and 1.2-fold (p < 0.05) higher in *hns*Δ93 than wild-type, respectively. In addition *aceA* had 1.4-fold higher expression (p = 0.07).

**Figure 3:**
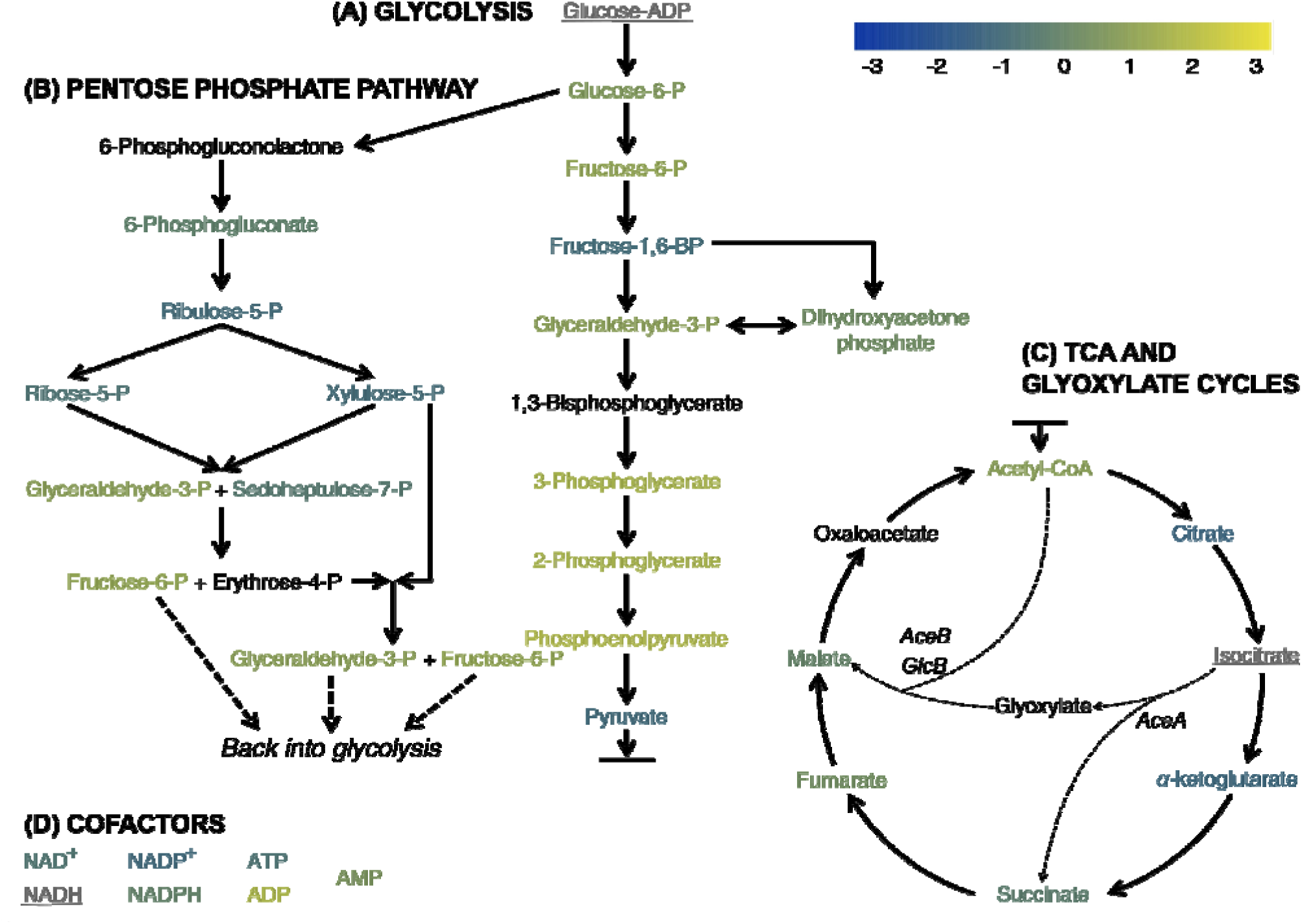
Overview of changes in central carbon metabolism in quiescent *E. coli* at 240 min. The color of each metabolite name corresponds to the average mean-centered fold change in concentration (n = 4) at the end of the time-course experiment for quiescent cultures (*hns*Δ93 with 3 mM indole). Metabolites in grey, underlined text had concentrations below the limit of detection. Those in black text were not possible to detect with our system. Italic font indicates the names of enzymes involved in the glyoxylate pathway.

These results lend some support to the hypothesis that the glyoxylate cycle is more active in the *hns*Δ93 mutant, although further work would be necessary to confirm whether the modest increases in gene expression have a significant effect on metabolic activity.

A key function of the TCA cycle is to reduce NAD^+^ to NADH, which acts as an electron donor for the electron transport chain. In our experiment, the concentration of NAD^+^ and the related molecule NADP^+^ slowly decreased over 4 h under all conditions, but remained highest for *hns*Δ93 with 3 mM indole (Supplementary Fig. 6). The concentrations of the reduced forms of both molecules were also lower than the oxidized forms; in the case of NADH, below the limit of detection. Furthermore, adenosine phosphate concentrations remained more constant in quiescent cultures than under other conditions. Therefore, the quiescent state must involve a homeostatic mechanism by which cofactor concentrations are controlled.

### 3.4 *Global protein expression changes are primarily due to the* hns *mutation*

Since H-NS regulates approximately 5% of all gene expression in *E. coli* (Hommais et al., 2001) we speculated that the *hns*Δ93 mutation might affect the expression of proteins involved in metabolism and stress response. Therefore, we drew samples for protein expression profiling in parallel with samples for metabolite analysis 1 h into the time-course experiment. 2-dimensional differential in-gel electrophoresis (2d-DIGE) showed that almost all the variation in protein expression was due to the strain genotype rather than indole addition (Fig. 4a,b). A total of 424 protein spots were present in every spot map (complete cases), with 32% of those differing significantly (p ≤ 0.05) due to the strain (Fig. 4c). Using more stringent criteria, we identified 43 protein spots with ≥ 3-fold increase or decrease in expression compared to the control (p ≤ 0.001). ANOVA for the 43 selected spots showed that 39 spots varied due to the strain and 2 spots due to indole addition. The remaining 2 spots varied due to a combination of indole and strain, although there was no interaction.

**Figure 4:**
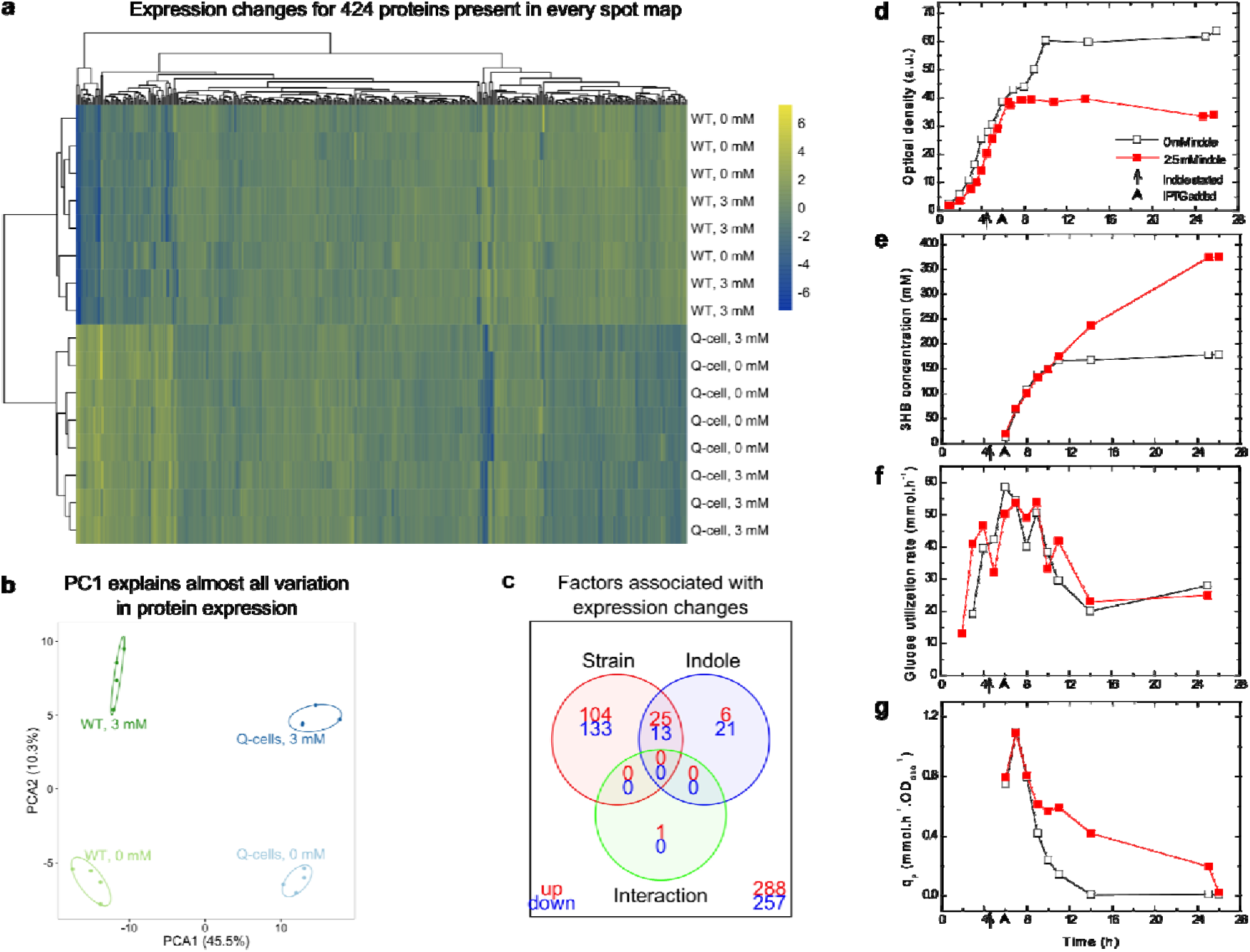
**Global changes in protein expression, and increased production of 3HB in quiescent cultures. a**, Heat map of fold-changes in protein expression determined by 2d-DIGE for 424 complete cases. **b**, Principal component analysis on the protein expression data confirms a single principal component accounted for the majority of variation, with separation primarily by strain. **c**, Significantly up-and down-regulated proteins from the complete cases, grouped by the cause(s) of the change in expression, as determined by ANOVA (adj. p < 0.05). **d -g**, Fed-batch cultures of *E. coli* W3110*hns*Δ93 producing 3HB, with or without the addition of indole (2.5 mM), showing that as cells enter quiescence their metabolic effort is diverted away from biomass production and can be harnessed for the production of 3HB by metabolic pathway engineering. We show here the results of the experiment in which 2.5 mM indole was used. Similar results were achieved with other indole concentrations but are not directly comparable. **d**, Indole addition (arrows) caused the culture to become quiescent within 2 h. **e**, Cumulative 3HB production following induction with IPTG (arrow heads). While the quiescent culture remained productive, the control stopped producing 3HB upon entry into stationary phase. **f**, Glucose utilization rates were the same for both strains throughout the experiment. **g**, **S**pecific 3HB formation rates (q_p_) were initially the same, but reduced in the control during stationary phase.

The 43 selected spots were excised from the gel, digested by trypsin and subjected to peptide fingerprinting analysis via mass spectrometry. The results were queried against the SwissProt database using MASCOT v.2.4.1, and we were able to identify 12 spots with a high certainty of representing a single protein (or, in one case, two) (Table 2). Of the 12 proteins identified, 8 are regulated by H-NS (www.ecocyc.org). Of these, 7 are involved in stress response functions and were up-regulated in the *hns*Δ93 strain. This suggests Q-cells might be ‘primed’ for metabolic stresses induced by indole addition to the medium. Interestingly, PyrB expression was also reduced in *hns*Δ93. We noted earlier that the *hns*Δ93 mutant grew more slowly than the wild-type in mineral salts medium. PyrB knockout mutants are unable to grow in M9 medium due to a reduction in nucleotide synthesis, so the reduction in PyrB expression in the mutant might explain this observation (Patrick et al., 2007).

**Table 2:** Identities and properties of proteins with greater than 3-fold change in concentration from the control (p < 0.001)

| Spot ID | Protein Name | Gene | Theoretical M<sub>r</sub> | Fold Change<sup>†</sup> | Change due to: | Pathway / function |
| --- | --- | --- | --- | --- | --- | --- |
| 877 | NADP-specific glutamate dehydrogenase | gdhA | 48,778 | -4.14 | Strain | Glutamate synthesis |
| 1319 | Aspartate carbamoyltransferase | pyrB | 34,463 | -6.23 | Strain | Pyrimidine nucleotide synthesis |
| 1932* | Protein YciF | yciF | 18,643 | 14.90 | Strain | Osmotic shock |
| 1988* | Aspartate carbamoyltransferase regulatory chain | pyrI | 17,338 | -4.59 | Strain | Regulatory partner of PyrB |
| 2090* | Protein YciE | yciE | 19,007 | 14.67 | Strain | Osmotic shock |
| 2050* | DNA protection during starvation protein | dps | 18,684 | 5.82 | Strain | DNA protection |
| 2121* | 50S ribosomal protein L10 | rlpJ | 17,757 | 6.94 | Strain | Ribosome component |
| 2145* | Uncharacterised protein YgaU | ygaU | 16,053 | 3.68 | Strain | Osmotic shock |
| 2153* | Peroxioredoxin | osmC | 15,193 | 4.70 | Strain | Oxidative stress |
| 2235 | 1) 50S Ribosomal protein L9 | 1) rplI | 1) 12,149 | 3.40 | Indole | 1) Ribosome component |
|  | 2) Outer membrane assembly factor | 2) bamE | 2) 12,408 |  |  | 2) Unknown |
| 2314 | Thioredoxin-1 | trxA | 11,913 | 3.25 | Strain | Control of redox potential |
| 2375* | Protein YjbJ | yjbJ | 8,320 | 8.90 | Strain | Osmotic shock |

\* Regulated by H-NS
<sup>†</sup> Average fold change in expression level (n = 4) 1 h into the time-course experiment. Positive numbers represent increased concentration and negative numbers represent a decrease.

### 3.5 Application of Q-cells to the production of 3-hydroxybutyrate

The build-up of glycolysis intermediates, and the fact that fewer resources are diverted towards growth, potentially make Q-cells an ideal cell factory for small molecule production. 3HB was chosen as a model for metabolite production as it requires the expression of only three heterologous enzymes for its production from Ac-CoA and its presence in the growth medium can be assayed simply with commercially-available kits. To produce 3HB, two molecules of Ac-CoA are condensed by a β-ketothiolase (PhaA from *Cupriavidus necator*) and reduced at the β-position by an acetoacetyl-CoA reductase (PhaB from *C. necator*), before the CoA moiety is removed by an overexpressed *E. coli* thioesterase B (TesB) (Liu et al., 2007; Tseng et al., 2009), as illustrated in Supplementary Fig. 7. We expected that this biosynthetic pathway would exploit the large accumulation of PEP and Ac-CoA in quiescent cultures (Fig. 1d and Table 1).

We introduced plasmid pTrctesBphaAB into *E. coli* W3110*hns*Δ93 to enable it to produce 3-hydroxybutyrate (3HB) in an IPTG-inducible manner. Terrific broth (TB) with glucose as the carbon source was chosen as the culture medium to avoid the slow growth of the *hns*Δ93 strain in M9GYT (Fig. 1a) (Tartof and Hobbs, 1987). Pilot experiments demonstrated that quiescence could be achieved with 2.5 mM indole so we used this concentration to minimize the potential for indole toxicity. After 26 h the quiescent culture reached an OD_600_ of 34.0, compared to 63.7 for a control culture with ethanol added in place of the indole solution (Fig. 4d). This corresponded to dried cell weights (DCW) of 13.3 g.L^-1^ and 24.8 g.L^-1^ respectively. Growth of the cultures was almost identical until indole addition, after which the Q-cell culture ceased growing almost immediately. Despite the reduced cell density, the 3HB concentration in the Q-cell culture reached 374.8 mM (39.4 g.L^-1^) at 26 h compared to 178.4 mM (18.6 g.L^-1^) for the control (Fig. 4e). Therefore, over the course of the experiment, the product yield (grams of product formed per liter divided by grams of cells formed per liter) of the quiescent culture was 4.0-fold greater than for the control (2.97 against 0.75), while the absolute concentration in the culture supernatant was 2.1-fold greater.

Importantly, the rates of substrate (glucose, S) utilization (dS/dt) were very similar for both cultures at all stages of the fermentation (Fig. 4f). On the other hand, the specific product (3HB, P) formation rates (given by 1/X(dP/dt), where X is biomass), representing the amount of 3HB produced per unit of time per unit of biomass, were initially similar but diverged 9 h into the experiment (Fig. 4g). Comparison with Fig. 4d demonstrates that the divergence corresponds with entry of the control culture into stationary phase. 3HB production is linked to the metabolic rate of the cell. Therefore, the control culture became unproductive as metabolism slowed upon entry into stationary phase, whereas the quiescent culture continued to produce 3HB. It is this extended production period that accounts for the superior productivity of quiescent cultures. Ceasing growth clearly allows quiescent cultures to take advantage of the large pool of available metabolites that were revealed by our metabolomics analysis.

## 4 Discussion

We show here that the *hns*Δ93 mutation leads to wide-scale changes in protein expression. These changes did not greatly affect the balance or concentration of metabolites between the strains. Most significantly, stress response genes were upregulated, in agreement with previous characterization of protein expression changes in an *hns* knockout strain (Laurent-Winter et al., 1997). A detailed survey of expression changes due to indole has not been carried out in *E. coli*, but it has been suggested that proteins involved in drug and acid resistance (Hirakawa et al., 2010, 2005; Lee et al., 2010), biofilm formation (Di Martino et al., 2003; Lee et al., 2007; Wood, 2009), amino acid metabolism (Wang et al., 2001), virulence (Hirakawa et al., 2009), and even inter-species communication (Lee and Lee, 2010; Vega et al., 2013) are all affected by indole. A microarray study of indole-induced expression changes in *Salmonella* found 77 differentially expressed proteins (Nikaido et al., 2012). Although we imposed more stringent cut-off parameters for the selection of significant expression changes, our results are in agreement that indole does not affect the expression of a large number of genes.

DNA replication and gene transcription in *E. coli* are coordinated throughout the growth cycle by controlling the topological structure of the chromosome (Balke and Gralla, 1987; Hsieh et al., 1991; Ohniwa et al., 2006). It has also been suggested that a high density of DNA gyrase binding sites (and consequently high superhelical density) near to the replication origin is correlated with genes that are expressed early in the growth cycle (Lau et al., 2004; Sobetzko et al., 2012). Indole inhibits DNA gyrase, but is not thought to have a strong effect at 3 mM (Field and Summers, 2012). Therefore, the protein expression and metabolic effects we observed are unlikely to be a consequence of DNA gyrase inhibition by indole. As noted before, other mutations in *hns* do not allow the induction of quiescence (Rowe and Summers, 1999). Therefore, we conclude that the *hns*Δ93 mutation affects the binding of H-NS to DNA in a way that loosens its control over the expression of a subset of the genes which it inhibits, and that this subset includes a large number of stress response genes.

It is not necessary for indole to affect protein expression in order to influence metabolite concentrations since metabolic balance is controlled largely through modulation of enzyme activity (Ishii et al., 2007). As a protonophore, indole reduces the PMF, leading to increased proton pumping by the electron transfer chain (Korshunov et al., 1997). This stimulates a demand for NADH from the TCA cycle, leading to an increased rate of respiration. However, in our experiments the TCA cycle was disrupted. Continued supply of glucose into glycolysis but reduced activity in the TCA cycle led to the build-up of glycolytic intermediates, particularly at the junction between glycolysis and the TCA cycle.

Our results suggest that the *hns*Δ93 mutation essentially ‘primes’ cells for the effect of indole on metabolism and prepares the cells for quiescence. When indole is present, metabolism proceeds in a redox-imbalanced fashion, resulting in a reduction in NAD^+^ and NADP^+^ concentrations as the pathways that regenerate them are inhibited (Fig. 3). In the wild-type strain this soon leads to oxidative damage by reactive oxygen species (ROS) (Cabiscol et al., 2000; Krapp et al., 2011). However, in *hns*Δ93, elevated concentrations of stress-response proteins may enable the ROS to be neutralized, partly by oxidizing NADH and NADPH. Therefore, the increased defense against oxidative stress in Q-cells may also act to regenerate cofactors, allowing for continued glucose metabolism. This model of indole action could shed further light on the metabolic shift that takes place as wild-type cells transition from exponential to stationary phase (Gaimster and Summers, 2015). For example, it might be of benefit for cells to repress the TCA cycle when less energy is required to power cell growth and division, and divert metabolic energy towards cell maintenance pathways leading from the PPP and glycolysis.

As an example of the productive capacity of the Q-cell system we demonstrated enhanced production of 3HB. Other than adjusting the indole concentration, we made no attempt to optimize the production conditions. However, the 39.4 g.L^-1^ of 3HB produced here was still greater than the highest previously reported productivity of 3HB of 12.2 g.L^-1^ from 16.7 g.L^-1^ DCW (Liu et al., 2007).

By better understanding the mechanism of entry into quiescence we have been able to identify promising starting points for metabolic engineering to improve the *hns*Δ93 strain for the production of commercially significant products through exploitation of the accumulated glycolysis intermediates. The uncoupling of production from biomass generation clearly leads to large increases in efficiency. Therefore, we believe that with additional metabolic engineering and optimization of growth conditions, the Q-cell system will become a powerful and flexible tool for industrial-scale production of a variety of high value chemicals.

## 5 Methods

### Strains and growth conditions

The wild-type strain used as the control was *E. coli* W3110 (ATCC 27325). To create the Q-cell strain, the *hns*Δ93 mutation was introduced to the wild-type strain using the lambda Red recombinase system, as described previously (Chen et al., 2015). Strains were stored with glycerol at −80 °C and streaked onto Luria-Bertani (LB) agar plates before use. A 5 ml starter culture was then grown in LB liquid medium for 8 h at 37 °C and used to inoculate further cultures for the experiments. The *hns*Δ93 mutation confers kanamycin resistance to the Q-cell strain. For selection purposes, Q-cell plates and starter cultures contained kanamycin (30 μg.ml^-1^). However, to keep the conditions used for each strain as similar as possible, no antibiotics were used for experimental cultures, except when necessary for maintenance of the 3-hydroxybutyrate production plasmid.

For the metabolome and proteome studies, cultures were grown in M9 medium (12.8 g.L^-1^ Na_2_HPO_4_.7H_2_O, 3 g.L^-1^ KH_2_PO_4_, 0.5 g.L^-1^ NaCl, 1 g.L^-1^ NH_4_Cl and 0.24 g.L^-1^ MgSO_4_) supplemented with 4 g.L^-1^ glucose, 2 g.L^-1^ yeast extract and 1 ml.L^-1^ of a trace elements solution (9.7 g.L^-1^ FeCl_3_, 7.8 g.L^-1^ CaCl_2_, 0.218 g.L^-1^ CoCl_2_.6H_2_O, 0.156 g.L^-1^ CuSO_4_.5H_2_O, 0.118 g.L^-1^ NiCl_3_.6H_2_O and 0.105 g.L^-1^ CrCl_3_.6H_2_O dissolved in 0.1 M HCl). Seed cultures (300 mL) were inoculated with 3 mL of LB starter culture in 1 L baffled Erlenmeyer flasks and grown for 16 h overnight with shaking at 300 rpm.

Following inoculation of the fermentor (final volume of 3 L in a 10 L vessel), cultures were grown in batch mode until the glucose supply was exhausted, indicated by a rise in dissolved oxygen (DO) and pH, and confirmed by testing with a hand-held FreeStyle Optium glucose meter (Abbott Laboratories, UK). Feed solution (180 g.L^-1^ glucose, 22 g L^-1^ NH_4_Cl, 5.42 g.L^-1^ MgSO_4_) was then introduced at 1.5 mL min^-1^ by peristaltic pump. Throughout the experiment, the temperature was controlled at 37 °C, agitation speed was a constant 500 rpm (sufficient to maintain DO above 40 %) and pH was maintained at 7.0 by addition of 5 M NaOH as necessary. Control of foaming was only necessary for wild-type cultures without indole, and was achieved by automatic addition of antifoam agent A (Sigma-Aldrich, US) controlled by a foam level probe.

After a 30 min equilibration period, indole dissolved in ethanol (1 M) was added in a single shot to a final concentration of 3 mM. An equivalent volume of pure ethanol was added for no-indole controls. The growth of each culture was monitored by measuring the optical density at 600 nm (OD_600_) of samples. W3110*hns*Δ93 cultures grew slower than the wild-type both before and after indole addition. Despite the different growth rates, the OD_600_ at the start of the feeding stage was the same for all cultures.

For the production of 3-hydroxybutyrate (3HB), a similar strategy was followed to that described above. However, a complex medium could be used as it was no longer necessary to avoid activating alternative metabolic pathways, as in the metabolome study. This also allowed both strains to grow at equal rates. We used terrific broth (TB) at all stages of the 3HB production experiments. TB medium consists of 24 g.L^-1^ yeast extract, 12 g.L^-1^ peptone, 9.4 g.L^-1^ K_2_HPO_4_, 2.2 g.L^-1^ KH_2_PO_4_, 4 g.L^-1^ glucose and 2.4 g.L^-1^ MgSO_4_. A 100 mL pre-culture of *E. coli* W3110*hns*Δ93/pTrctesBphaAB was grown overnight in TB medium at 37 °C then used to inoculate the fermentor (5 L vessel), which contained a further 1.9 L of fresh TB medium. Ampicillin (100 μg/mL) was included in all growth media and the nutrient feed, to select for cells containing the plasmid.

When the original glucose supply was depleted (signaled by a rise in pH), a feed medium consisting of 190 g.L^-1^ glucose, 108 g.L^-1^ peptone, 84 g.L^-1^ yeast extract, 9.4 g.L^-1^ K_2_HPO_4_, 2.2 g.L^-1^ KH_2_PO_4_ and 1.2 g.L^-1^ MgSO_4_ was pumped into the fermentor at an initial flow rate of 0.28 mL.min^-1^. The ratio of glucose to complex nitrogen in this feed was kept much higher than TB medium to provide a better stoichiometric balance. The flow rate was increased exponentially so as to double every 2 h pre-induction to keep pace with the growth rate of the cells. Indole was dissolved in ethanol to 150 mM and pumped into the fermentor at a rate of 0.28 mL.min^-1^ for 2 h, through a tube that exited below the level of the medium to ensure good dissolution. Induction was done with IPTG (1 mM final concentration) 30 min after the start of the indole feed. The feed rate post induction was kept constant since we observed a drop in the growth rate of the cells.

Cell growth was measured as OD_600_ following appropriate dilution and as cell dry weight (CDW) by collecting the cells from two 1 mL samples through centrifugation and drying in an oven until constant weight. The average of the two weights was reported. Glucose concentrations were recorded using a medical glucose meter and disposable enzyme assay strips. substrate consumption rates (with respect to glucose) were calculated from a material balance on the amount fed per unit time and the residual glucose concentration in the culture medium. Following centrifugation at 17,000 × g for 2 minutes to remove the cells, samples of the supernatant (1 mL) were kept at −80 °C before testing for 3HB concentrations using a β-hydroxybutyrate colorimetric assay kit (Cayman Chemicals, US).

### 3-hydroxybutyrate production plasmid construction

Plasmid pTrctesBphaAB was constructed from plasmid pTrcphaCAB_Re_ which was generously provided by Dr. Takeharu Tsuge (Tokyo Institute of Technology, Japan) (Kahar et al., 2005), by replacing the *phaC* gene with *tesB*. The artificial operon thus created encoded for a 3-enzyme pathway in which 3-hydroxybutyryl-CoA is produced from acetyl-CoA by PhaA and PhaB, and the CoA moiety is then removed by TesB to leave 3-HB. The *tesB* gene was amplified by PCR from plasmid pCA24N:TesB (Kitagawa et al., 2005) and the product was purified using a QIAQuick PCR purification column (Qiagen, USA). Plasmid pTrcphaCAB_Re_ was linearized by digestion with EcoRI and SalI and the plasmid backbone was then ligated with the *tesB* fragment by Gibson Assembly according to the manufacturer’s instructions of a kit provided by New England Biolabs (US). The primers used for Gibson Assembly were:

Forward: AACAATTTCACACAGGAAACAGACCATGGAATTCAGATCTTTCG

AATAGTGACGGCAGAGAGACAATCAAATCATG

Reverse: CAAAACAGCCAAGCTTGCATGCCTGCAGGTCGACTCTAGAGGAT

CCAAACCCGGTGAATTGGCGCA

### Time-course study of metabolomics changes

Samples were taken immediately before indole addition and at six other points over the next 4 hours (5, 15, 30, 60, 120 and 240 min after indole addition). Approximately 20 mL samples were drawn aseptically from the fermentor, 1 mL aliquots of which were used for the metabolomics study. The OD_600_ of each sample was measured with appropriate dilution and recorded, and later used to normalize the metabolite concentrations. We obtained samples for metabolite quantification by vacuum filtering 1 mL of each culture and quenching the cells by immersion in cold (-80 °C) methanol within 30 sec of withdrawal. The samples were stored in methanol at −80 °C until preparation for analysis.

### LC-MS/MS analysis of metabolites

Prior to analysis, metabolites were extracted using a previously described method with modifications (Soga, 2007; Yoshida et al., 2008). Briefly, the metabolites were extracted into a 1.2Lml solvent mixture (CHCl_3_:H_2_O, 1:1, v/v) containing 10 μg.L^−1^ of D-(+)-camphor-10-sulfonic acid as an internal standard for semi-quantitative analysis. After centrifugation at 15,000 ×Lg at 4 °C for 5 min, 10 μL of the upper phase was used for quantification of intracellular metabolites by high-performance liquid chromatography coupled with electrospray ionization tandem mass spectrometry (LCMS-8040 triple quadrupole LC-MS/MS spectrometer; Shimadzu, Japan) as described previously (Luo et al., 2007).

### GC-MS analysis of metabolites

For GC-MS analysis, 70 μL of the upper phase, as for LC-MS/MS analysis, was transferred to a new tube and vacuum dried. The dried residue was derivatized for 90 min at 30 °C in 20 mg.mL^-1^ methoxyamine hydrochloride in pyridine (20 μL). subsequently, trimethylsilylation (TMS derivatization) was performed for 30 min at 37 °C and then for 2 h at room temperature with N-methyl-N-(trimethylsilyl)trifluoroacetamide (MSTFA, 50 μL) (Roessner et al., 2000; Strelkov et al., 2004). GC-MS was carried out using a GCMS-QP2010 Ultra (Shimadzu, Japan) equipped with a CP-Sil 8 CB-MS capillary column (30 m × 0.25 mm × 0.25 μm; Agilent, USA). Helium was used as the carrier gas with a flow rate of 2.1 mL.min^-1^. The injection volume was 1 μL with a split ratio of 1:10. An initial oven temperature of 60°C was maintained for 10 min, then raised to 315 °C at 15 °C.min^-1^, and maintained for 6 min. The total running time was 33 min. The other settings were as follows: 250 °C interface temperature, 200 °C ion source temperature, and electron impact (EI) ionization at 70 eV.

### HPLC analysis for measurement of organic acids in medium

Supernatant of the cell broth, recovered after centrifugation, was used for HPLC using a Prominence HPLC System (Shimadzu, Japan) with a conductivity detector and two Shim-pack SCR-102H columns (300 mm x 8.0 mm; I.D., 7 μm; Shimadzu, Japan). The column temperature was 48 °C and the flow rate of the mobile phase (5 mM p-toluenesulfonic acid; p-TSA) was 0.8 mL.min^-1^. The flow rate of the pH buffering solution for the detector (5 mM p-TSA, 20 mM Bis-Tris, and 0.1 mM EDTA-4H) was 0.5 mL.min^-1^.

### 2d-DIGE comparison of protein expression

The same 20 mL samples from the fermentation cultures as described for the metabolomics study were used for the proteomic study. A 10 mL aliquot of each sample was centrifuged at 4000 × g for 15 min to recover the cells. The pellet was then washed 3 times by repeated suspension and centrifugation, in an ice-cold buffer consisting of 10 mM Tris (pH 8.0) and 5 mM magnesium acetate. Washed pellets were stored at −80 °C until all samples had been collected, and then prepared for analysis simultaneously.

Cells were resuspended in lysis buffer (7 M urea, 2 M thiourea, 30 mM Tris and 4% (w/v) CHAPS) and lysed by sonication on ice. The sonication protocol consisted of 12 cycles of 10 sec with 10 sec rest periods between. The pulse amplitude was set to 15 Amp resulting in a pulse power of 8 – 9 W. After complete lysis, the protein concentration was measured using a Bradford microplate assay procedure. Controls consisting of bovine serum albumin were prepared in the same lysis buffer to prepare a standard curve. The pH of each sample was also checked and found to be close to pH 8.5, which is optimal for the labelling procedure.

An internal standard was created by pooling equal amounts of protein from every sample. For the test samples, 50 μg of protein was labelled for each sample. Labelling was carried out with the CyDye DIGE Fluor minimal labelling kit (GE Healthcare, US) using Cy2 for the internal standard and Cy3 and Cy5 for test samples. Groups of two test samples were then combined together with an internal standard sample, and mixed with an equal volume of 2× sample buffer (7M urea, 2 M thiourea, 2% (w/v) CHAPS, 0.5% IPG buffer (pH 3 – 11 NL) and De-streak reagent). Finally, the volume was made up to 450 μL with rehydration solution (7 M urea, 2 M thiourea, 2% (w/v) CHAPS). Each sample was loaded onto a 24 cm Immobiline DryStrip (pH 3 – 11 NL) using the rehydration method, with rehydration proceeding for 12 h at 20 °C. Isoelectric focusing then proceeded with an initial step of 500 mV for 1 h, followed by a gradient to 100 mV over 8 h, a gradient to 8000 mV over 3 h and a final step with the voltage held at 8000 mV for 3.75 h.

The second dimension electrophoresis was conducted using the Ettan DALT gel and electrophoresis system (GE Healthcare, USA). 24 cm pre-casted gels were used following equilibration of the focused IEF strips. 8 gels were run simultaneously with 12 mA per gel, with a recirculating pump and chiller to maintain the buffer temperature at 15 °C, for 17 h until the bromophenol blue dye front just reached the end of the gel. The gels were then immediately scanned using a Typhoon 9400 variable mode imager (Amersham Biosciences, UK) and following the manufacturer’s recommended settings. The images were analyzed to detect changes in protein concentrations using DeCyder 2-D v.6.5 image analysis software.

### Protein digestion and identification

A preparatory gel containing 500 μg of unlabeled, pooled samples was run under the same conditions as for the analytical gel, and post-stained using SYPRO Ruby (Fisher Scientific, Japan). Following spot matching in DeCyder 2-D, the selected spots were excised using an Ettan spot picker (Amersham Biosciences, UK). The spots were de-stained in 50 mM ammonium bicarbonate containing 50%acetonitrile (50 μL) for 10 min at 37 °C, dehydrated with acetonitrile (25 μL) then dried in a vacuum centrifuge. To reduce cysteine residues, 100 mM ammonium bicarbonate with 10 mM dithiothreitol (25 μL) was added for 15 min at 50 °C, then 250 mM iodoacetamide in 100 mM ammonium bicarbonate (2 μL) was then added and the spots were incubated for 15 min at room temperature in the dark for alkylation. After washing and dehydration as before, the protein in the dried gel debris was digested at 37 °C overnight with 100 ng/10 μL modified trypsin solution. The digested protein fragments were collected from the supernatant and extracted from the gel debris by the addition of 50-80% acetonitrile containing 1% trifluoroacetic acid (3 × 25 μL).

The resulting protein sample was resolved in 2% acetonitrile containing 0.1% trifluoroacetic acid and applied to the liquid chromatography (LC) system (Advance nanoLC; Bruker-Michrom, USA) coupled to an LTQ linear ion trap mass spectrometer (ThermoFisher, USA) with a nanospray ion source in positive mode. The peptides were separated on a NANO-HPLC C18 capillary column (0.075 mm ID × 150 mm length, 3 mm particle size, Nikkyo Technos, Japan) using a linear gradient (25 min, 5-35% acetonitrile containing 0.1% formic acid) at a flow rate of 300 nL/min. The LTQ-MS was operated in top-3 data-dependent scan mode. The precursor ions were selected automatically for MS/MS analysis on the basis of their signal intensities. The parameters of LTQ were as follows: spray voltage, 2.3 kV; capillary temperature, 250 °C; mass range (m/z), 400-1800; collision energy, 35%. Raw data was acquired by Xcalibur software. The MS/MS data were searched against the SwissProt 2014_10 database using MASCOT v.2.4.1 software (Matrix Science, UK). The MASCOT search parameters were as follows: enzyme, trypsin; fixed modifications, carbamidomethyl (Cys); variable modifications, oxidation (Met); peptide mass tolerance, ± 1.5 Da; fragment mass tolerance, ± 0.8 Da; max. missed cleavages, 1. Significant MASCOT scores were defined with p ≤ 0.05.

## Acknowledgments

We would like to thank Dr Kenji Ohtawa (Riken Support Unit for Biomaterial Analysis, Japan) for technical assistance with 2D-DIGE, Dr Masaya Usui (Riken Support Unit for Biomaterial Analysis, Japan) for technical assistance with identification of protein spots, Dr Matthew Davey (Plant Sciences Department, University of Cambridge, UK) for helpful discussions about data analysis and Dr Antonio De León Rodríguez (IPICYT, Mexico) for critical reading of the manuscript. This project was funded by the Riken Foreign Postdoctoral Researcher program.

